# A comparison of data integration methods for single-cell RNA sequencing of cancer samples

**DOI:** 10.1101/2021.08.04.453579

**Authors:** Laura M. Richards, Mazdak Riverin, Suluxan Mohanraj, Shamini Ayyadhury, Danielle C. Croucher, J. Javier Díaz-Mejía, Fiona J. Coutinho, Peter B. Dirks, Trevor J. Pugh

## Abstract

Tumours are routinely profiled with single-cell RNA sequencing (scRNA-seq) to characterize their diverse cellular ecosystems of malignant, immune, and stromal cell types. When combining data from multiple samples or studies, batch-specific technical variation can confound biological signals. However, scRNA-seq batch integration methods are often not designed for, or benchmarked, on datasets containing cancer cells. Here, we compare 5 data integration tools applied to 171,206 cells from 5 tumour scRNA-seq datasets. Based on our results, STACAS and fastMNN are the most suitable methods for integrating tumour datasets, demonstrating robust batch effect correction while preserving relevant biological variability in the malignant compartment. This comparison provides a framework for evaluating how well single-cell integration methods correct for technical variability while preserving biological heterogeneity of malignant and non-malignant cell populations.

## BACKGROUND

Single cell transcriptome profiling of tumours has the ability to decode cellular heterogeneity in both the malignant and microenvironmental compartments. While profiling individual samples will incrementally expand our understanding of the cell types and states present within a single tumour, integrating data across different patients, longitudinal timepoints and cancer types is crucial to uncovering shared cell states underlying disease pathogenesis and treatment resistance. However, biological variation can easily be masked by technical factors introduced during sample processing and sequencing^1–3^, making it difficult to batch correct and integrate data from multiple sources.

Previous single cell batch correction benchmarking studies evaluated algorithm performance on simulated data or datasets derived from healthy tissues and peripheral blood mononuclear cells^4–6^. Despite an abundance of data, no single cell batch-effect correction methods are designed for or benchmarked on single cell datasets containing malignant cells^7^. Due to the inherent biological complexity both within and between tumours, these samples present unique technical challenges for batch correction that are not represented in previous benchmarking efforts. Unlike their normal counterparts, malignant cancer cells often express patient-specific transcriptional programs driven by somatic alterations at the DNA level, resulting in clustering patterns with little overlap between tumors from different individuals^8–13^. Additionally, in some cancer types, malignant cells hijack developmental pathways and share transcriptional programs with their normal cellular counterparts^14–17^. Transcriptional similarity between malignant and normal cells poses another batch correction challenge, as over-integration could result in co-clustering of fundamentally different cell types, thereby masking underlying biological signals within these populations.

Here, we perform a benchmark of 5 data integration tools on 5 cancer single cell or single nuclei RNA-sequencing datasets. We further discuss a framework to evaluate the accuracy of an integration method in terms of limiting technical variability while preserving true biological heterogeneity between cancer patients in both malignant and non-malignant compartments of the tumour microenvironment. With the continued growth of tumour single cell transcriptomics, effective batch correction and data integration will be crucial to mitigate technical artifacts between studies to generate large-scale, pan-cancer cellular atlases.

## RESULTS & DISCUSSION

We evaluated the performance of 5 batch correction algorithms for their ability to integrate batches (separate single cell encapsulation experiments) while maintaining separation between dissimilar cell types and preserving integrity of malignant cell clusters that may have patient-specific transcriptional programs driven by somatic alterations^8–13^. We focused on data integration tools available in the R programming language, capable of integrating with the single cell genomics analysis package Seurat^18,19^. Seurat is widely used by cancer genomics researchers, and so compatible data integration tools will likely have high uptake by the community. Specifically, we compared uncorrected data to integrations generated with Conos^20^, fastMNN^21,22^, Harmony^23^, STACAS^24,25^ and the Seurat implementation of reciprocal principal component analysis^26^ (RPCA). We did not include Seurat’s canonical correlation analysis^19^ (CCA), as it employs a similar methodology to RPCA but has inferior performance on datasets that share only a subset of cell types, as often encountered when comparing cancerous tissues^5^. The underlying methodology of each tool is described in the **Methods.**

Batch correction algorithms were tested on 5 datasets (4 published, 1 newly generated) covering a diverse spectrum of cell and cancer types. Datasets contained mixtures of malignant and non-malignant cells found in the tumour microenvironment (ie. immune and stromal) in glioblastoma (GBM; “Richards-GBM-LGG”), oligoastrocytoma (LGG; “Richards-GBM-LGG”) **(Table S1)**, basal cell carcinoma^27^ (BCC; “Yost-BCC”), childhood acute lymphoblastic leukemia^28^ (cALL; “Caron-ALL”), hepatocellular carcinoma^29^ (HCC; “Ma-LIHC”) and clear cell renal cell carcinoma^30^ (cRCC; “Bi-RCC”). All datasets were derived from human samples and profiled with 10x Genomics single cell RNA-seq technologies with either live cells or nuclei as input. Collectively, these 5 datasets profiled the transcriptomes of 171,206 cells or nuclei from 68 tumour samples derived from 52 patients across 6 cancer types **(Figure S1A)**. We generated two sets of biological replicates from in our in-house glioma snRNA-seq dataset, providing a unique opportunity to better assess technical influences while limiting confounding patient-to-patient heterogeneity. We define a biological replicate as two independent single cell encapsulation experiments on two pieces of the same tumor (Tumour 1=B_P_GBM593.1, B_P_GBM593.2; Tumour 2=C_P_GBM577.1, C_P_GBM577.2), acknowledging that these replicates may exhibit intratumoural heterogeneity. Additionally, Richards-GBM-LGG and Yost-BCC contain longitudinal samples from the same patient before and after treatment. Before data integration, datasets were processed in a harmonized manner including data normalization, scaling and identification of variable genes **(Methods)**. Unlike simulated data or data from normal tissues, batches (samples) did not necessarily have similar cell type composition. Both cell type representation and proportion varied between samples and biological replicates within datasets **(Figure S1C)**.

To qualitatively evaluate integration results, we used Uniform Manifold Approximation and Projection (UMAP) visualizations before (uncorrected) and after data integration **(Figures 1**, **S2** **&** **S3**). To quantitatively assess batch correction, we computed the Local Inverse Simpson’s Index (LISI) metric^23^ using UMAP or LargeVis cell embeddings **(Figure 2)**. LISI has been shown to effectively assess whether groups of cells are well mixed across batches, and performs well amongst other batch assessment metrics (ie. adjusted Rand Index, average silhouette width, *k*-nearest neighbor batch-effect test^31^) on single-cell RNA-sequencing data^4–6^. For every dataset-integration method combination, we calculated the median scaled LISI metric to assess batch (sample) mixing within individual cell types and across entire datasets **(Figure 2**; **Methods)**. To permit comparison of LISI scores between datasets, we scaled scores between 0 and 1 using batch size **(Methods)**, where 1 represents a good integration defined by high mixing between batches.

**Figure 1.**
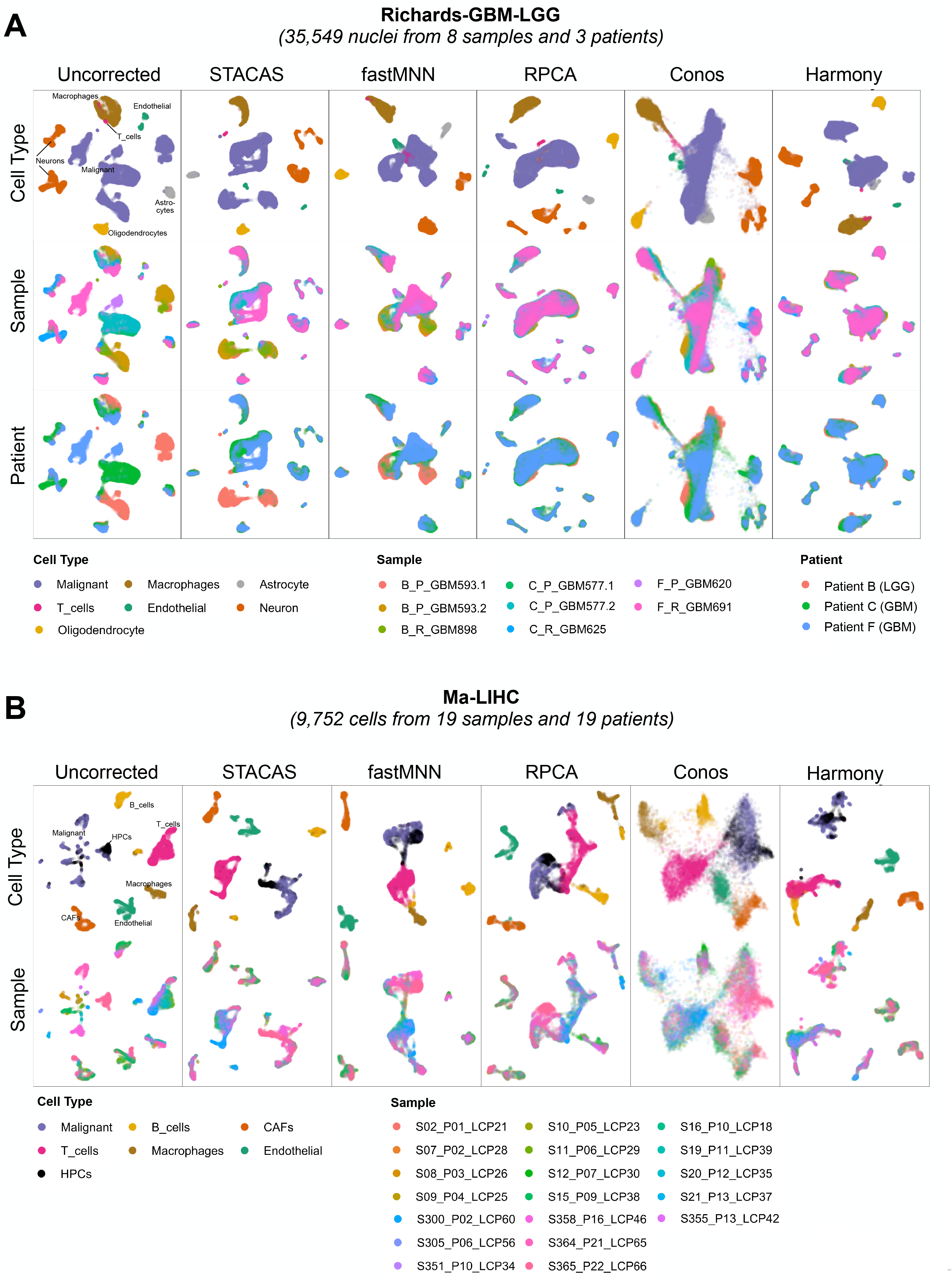
Qualitative comparison of data integration methods across cancer datasets. UMAP or LargeVis (Conos) visualizations of cells from **(A)** high-grade glioblastoma (GBM) and low-grade oligoastrocytoma (LGG) (Richards-GBM-LGG; n=35,549 cells), or **(B)** liver cancer (Ma-LIHC, n=9,752 cells). Each dot represents a cell. Cells are colored by cell type (top row), sample (middle row) or patient (bottom row). Columns represent either uncorrected (first column) or integrated data (columns 2 to 6). HPCs, hepatic precursor cells; CAFs, cancer associated fibroblasts.

**Figure 2.**
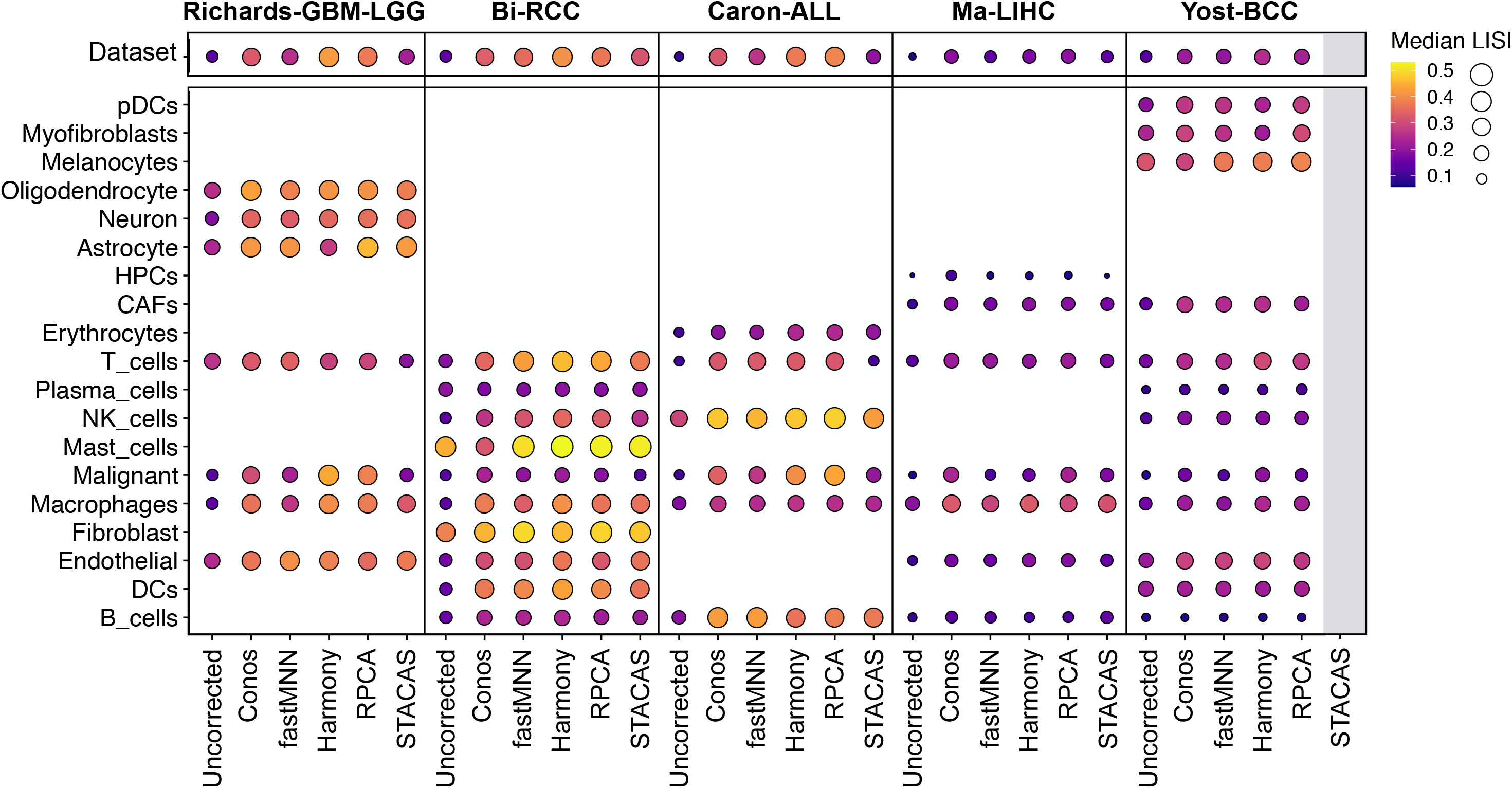
Evaluating data integration performance with Local Inverse Simpson’s Index (LISI) metric. Dot plot depicting sample mixing with scaled median LISI scores across the entire dataset (top panel) or within individual cell types (bottom panel) for each data integration method (x-axis). Color and size of dot represent median LISI score. The absence of a dot represents cell types that are not found in a given dataset. Grey bar denotes missing integration (unable to generate STACAS integration for Yost-BCC). Vertical black lines separate datasets. pDCs, plasmacytoid dendritic cells. DCs, dendritic cells; HPCs, hepatic precursor cells; CAFs, cancer associated fibroblasts.

First, we evaluated the ability for data integration methods to improve mixing between batches (samples) compared to the uncorrected analysis. Comparing UMAP visualizations, we observed a universal increase in sample mixing post-data integration across cellular compartments and cancer types **(Figure 1**, **S2** **and** **S3)**. In agreement with the visualizations, all data integration methods had higher dataset LISI scores compared to uncorrected data, demonstrating improved cell mixing between samples post-integration regardless of the method (Figure 2 **and** **S4A)**. Most notably, malignant cells clustered almost entirely by patient in the uncorrected analysis, but displayed increased overlap after integration. However, there are examples when increasing mixing between batches may not be biologically appropriate when analyzing tumour-derived data. For example, in Richards-GBM-LGG there are two different types of gliomas present - samples from high-grade glioblastoma (Patients C & F; GBM) and low-grade oligoastrocytoma (Patient B; LGG). Single cell profiling of these tumour types has shown they harbour different malignant transcriptional programs^10,15,32^, and so one would expect tumour cells from these two pathologies to have limited overlap. In our glioma cohort, clinical annotations revealed differences in the treatment regimens of GBMs and LGGs, as well as variable IDH1 mutation status across patients (IDH1 p.R100Q in F_P_GBM620 & F_R_GBM691) **(Table S1)**. Treatment history and somatic mutations may induce additional layers of transcriptional heterogeneity between the two glioma subtypes.

STACAS maintained the best separation between the two glioma types, while fastMNN maintained moderate separation. High-grade and low-grade gliomas almost completely overlapped after integration with Seurat’s-RPCA, Conos and Harmony–masking potential biological signal in the malignant compartment **(Figure 1A).** Similar trends were observed in the malignant compartments of kidney cancer (Bi-RCC), where patient-specific clustering patterns are visible in the malignant compartment with STACAS and fastMNN, without sacrificing improved overlap between batches in normal cell types **(Figure 1B)**.

Next, we compared batch mixing within non-malignant cell compartments. Across each uncorrected dataset, there are UMAP regions within normal cell types dominated by cells from individual samples, suggesting the presence of technical batch effects between samples **(Figure1**, **S2** **and** **S3).** Batch mixing improved in each normal cell type across datasets after-integration, regardless of method, with the exception of T cells in Richards-GBM-LGG integrated with STACAS (median LISI uncorrected = 0.26 vs. STACAS = 0.19) **(Figure 2)**. Further investigation is needed to determine if there are sample-specific T cell states present in these cancer types that could explain reduced sample mixing between batches. Comparing median cell type LISI scores of all normal cell types, Harmony had the highest batch mixing across all datasets **(Figure S4B)**, in concordance with results on normal tissues^4^. STACAS had the lowest batch correction but was still comparable to other integration methods and was an improvement over uncorrected data.

Finally, we assessed separation between dissimilar cell types. A good integration should eliminate batch-specific variability while limiting overlap between different cell types. Across cancer types and datasets, Conos integrations did not substantially separate different cell types. Dissimilar cell types frequently co-localized on LargeVis plots, such as endothelial, T cells and malignant cells in Richards-GBM-LGG and myofibroblasts and melanocytes in Yost-BCC (Figure 1A **and** **S3**). STACAS routinely provided good separation between cell types, while fastMNN, RPCA and Harmony had variable results depending on the dataset. STACAS and RPCA were the only algorithms able to cleanly separate a very small population of T cells in Richards-GBM-LGG, suggesting these tools may be beneficial when working with datasets containing rare subpopulations of cells specific to a subset of samples (**Figure 1A**). Surprisingly, all data integration methods, but not uncorrected data, struggled to separate macrophages and malignant leukemia cells in Caron-ALL, likely because of transcriptional similarity between monocytes/macrophages and differentiated monocyte-like leukemia cells^17^ **(Figure S2A)**. Differentiating cancer cells from their cell of origin, which may co-exist at high frequencies in the tumour microenvironment, represents a unique challenge when integrating cancer datasets and suggests that different cancer types may require different integration strategies to obtain optimal results.

Taken together, we identified STACAS as the best data integration algorithm based on its ability to eliminate technical biases and improve mixing of non-malignant cellular compartments across samples, while maintaining logical malignant clustering patterns representative of cancer pathology. Although STACAS was the top performer, it required extensive parameter optimization to integrate datasets with small batches (low cell counts less than 500) and had the longest run time **(Figure S4).** In these scenarios, the user can optimize the dist.pct and k.weight parameters during the anchor filtering and data integration steps, respectively, or remove problematic samples with few cells from the dataset entirely. However, this optimization is not always successful. For example, we were unable to generate a STACAS integration for Yost-BCC, likely due to the presence of several samples with low cell counts and high variability in batch (sample) size within the dataset (108 vs. 10,429 cells) (**Figure S1B** **and** **S3**; **Methods)**. Accordingly, we nominated fastMNN as a suitable alternative for integrating malignant datasets. fastMNN runs quickly (~7x faster than STACAS) without the need for parameter optimization and produces biologically reasonable integrations across cancer types. In our hands, Harmony and Conos often overintegrationed datasets, as demonstrated by inflated mixing between batches in the malignant compartment and poor separation between dissimilar cell types. In these cases, an uncorrected analysis may be more suitable than implementing a data integration tool.

## CONCLUSIONS

Single cell or nuclei transcriptomics datasets generated using different sequencing technologies or experimental conditions, such as dissociation protocols, will have batch-specific variation that needs to be accounted for during analysis. This benchmark addresses a gap on how well these methods integrate scRNA-seq data generated from tumor specimens. We demonstrated variable success of data integration methods across cancer types, highlighting the nuances of batch-correcting tumour scRNA-seq data and the importance of manual inspection to ensure integrations are biologically reasonable before proceeding with downstream analysis. We anticipate effective batch correction of cancer scRNA-seq datasets will enable the assembly of large-scale tumour atlases, revealing shared therapeutically-relevant cell types and states across cancer types.

## METHODS

### Single-nuclei RNA sequencing of gliomas

#### Generation of single nuclei suspensions

For the dataset “Richards-GBM-LGG”, single nuclei RNA-sequencing of gliomas was performed as previously described^11^. In brief, nuclei suspensions were generated from snap-frozen tumors. Tissues were minced on dry ice and dissolved in a lysis buffer, followed by homogenization with a pellet pestle. Nuclei integrity and quantity was assessed with SYBR Green II RNA Gel stain (Thermo Fisher Scientific). Nuclei were filtered through a 40-μm cell strainer and sorted for intact nuclei using DAPI (Sigma-Aldrich) on a BD Influx FACS sorter. Nuclei were re-suspended according to 10x Genomics concentration guidelines to obtain a target of 6,000 nuclei per sample.

#### Library preparation and sequencing

Library preparation was carried out as per the 10x Genomics Chromium single-cell protocol using the v2 chemistry reagent kit for 3’ expression profiling as previously described^11^. Libraries were sequenced on an Illumina HiSeq 2500 in High Output mode using the 10x Genomics recommended sequencing parameters to achieve the desired median read depth per cell (target mean 60,000 reads per nuclei).

#### Data pre-processing

We used the 10x Genomics CellRanger software pipeline (v.2.0) to demultiplex cell barcodes and map reads to a custom GRCh38 human reference transcriptome that included intron sequences to accurately quantify nuclear unspliced messenger RNA. We calculated the number of reads per cell barcode using the BamTagHistogram function in the Drop-seq Alignment Cookbook^33^ and subsequently determined the number of nuclei per library using the cumulative fraction of reads corresponding to cell barcodes. Cell barcodes were sorted in decreasing order and the inflection point was identified using the R package Dropbead^34^ (v.0.3.1) to distinguish between empty droplets and droplets containing a nucleus. The raw matrix of gene counts versus cells from CellRanger (v.2) output was filtered by the list of true cell barcodes from Dropbead. We processed the resultant unique molecular identifier (UMI) count matrix using the R package Seurat^18,19,26^ (v.4.0.1) as described below.

#### Quality control and data filtration

Following recent guidelines^2^ for frozen tumours profiled as single nuclei with v2 chemistry from 10x Genomics, we retained high-quality nuclei with at least 250 genes and 500 UMIs detected. We filtered out nuclei where >15% of UMIs came from mitochondrial genes. We removed lowly expressed genes present in 0.01% of cells in the average cell size. We predicted and removed potential doublets using Scrublet^35^ (v.0.2) with an expected_doublet_rate=0.06.

### Public dataset curation

For this study, we focussed on cancer scRNA-seq datasets that contained both malignant tumor cells and non-malignant cells from the tumour microenvironment. Only studies with raw UMI count matrices were considered. We did not perform additional cell-based quality control on count matrices, as this was already completed in the original publications. A detailed overview of all datasets can be found in **Figure S1**.

#### Ma-LIHC

Raw UMI count matrices and metadata, including cell type annotations, were downloaded from the Gene Expression Omnibus portal (GSE125449; https://www.ncbi.nlm.nih.gov/geo/query/acc.cgi?acc=GSE125449)^29^. We merged matrices from Set1 (n=12/19 tumours) and Set2 (n=7/12 tumours) for downstream analysis. To minimize ambiguity during integration, we removed cells labelled as “unclassified” in Ma-LIHC (n=194/9,946 cells). We relabelled a subset of cell type annotations provided by the authors to harmonize labels across datasets used in this study: “TAM” to “Macrophages”; “TEC” to “Endothelial; ‘HPC-like” to “HPCs”; “Malignant cell” to “Malignant”; “T cell” to “T_cells”; “B cell” to “B_cells”; “CAF” to “CAFs”.

#### Yost-BCC

Raw UMI count matrices and metadata, including cell type annotations, were downloaded from the Gene Expression Omnibus portal (GSE123813; https://www.ncbi.nlm.nih.gov/geo/query/acc.cgi?acc=GSE123813)^27^. We relabelled a subset of cell type annotations provided by the authors to harmonize labels across datasets used in this study: “CD4_T_cells”, “CD8_act_T_cells”, “CD8_ex_T_cells”, “CD8_mem_T_cells”, “Tcell_prolif”, “Tregs” to “T_cells”; “B_cells_1”, “B_cells_2” to “B_cells”; “Tumor_1”, “Tumor_2” to “Malignant”.

#### Caron-ALL

Raw UMI count matrices were downloaded from the Gene Expression Omnibus portal (GSE132509; https://www.ncbi.nlm.nih.gov/geo/query/acc.cgi?acc=GSE132509)^28^. Sample and cell-level metadata, including cell type annotations, were retrieved from the authors. Count matrices were merged across samples and used for downstream analysis. We relabelled some author cell type annotations as follows to harmonize labels across datasets used in this study: “ETV6.RUNX1.1”, “ETV6.RUNX1.2”, “ETV6.RUNX1.3”, “ETV6.RUNX1.4”, “HHD.1”, “HHD.2”, “PRE-T.1”, “PRE-T.2” to “Malignant”. We further subdivided author cell annotations “B cells + Mono” to “B_cells” and “Macrophages”, and “T cells + NK” to “T_cells” and “NK_cells”. Data was clustered as described below and clusters were annotated using expression of canonical cell type markers: B_cells (*MS4A1, BANK1*), Macrophages (*CD14, LYZ, FCGR3A, MS4A7*), NK_cells (*NKG7, GNLY*) and T_cells (*CD2, CD3G, CD4, CD8A*).

#### Bi-RCC

Raw UMI count matrices were downloaded from the Broad Institute Single Cell Portal (https://singlecell.broadinstitute.org/single_cell/study/SCP1288/)^30^. To minimize ambiguity during integration, we removed cells labelled as “Misc/Undetermined” in Bi-RCC (n=278/34,326 cells). We relabelled a subset of cell type annotations provided by the authors to harmonize labels across datasets used in this study: “41BB-Hi CD8+ T cell”, “MitoHigh T-Helper”, “41BB-Lo CD8+ T cell”, “T-Reg”, “Effector T-Helper”, “MitoHigh CD8+ T cell”, “Cycling CD8+ T cell”, “MX1-Hi CD8+ T cell”, “Memory T-Helper”, “NKT” to “T_cells”; “GPNMB-Hi TAM”, “FOLR2-Hi TAM”, “LowLibSize Macrophage”, “VSIR-Hi TAM”, “MitoHigh Myeloid”, “CXCL10-Hi TAM”, “CD16+ Monocyte”, “CD16- Monocyte”, “Cycling TAM” to “Macrophages”; “B cell” to “B_cells”; “MitoHigh NK”, “FGFBP2- NK”, “FGFBP2+ NK”, to “NK_cells”; “TP1”, “TP2”, “Cycling Tumor” to “Malignant”; “CD1C+ DC”, “CLEC9A+ DC” to “DCs”; “Plasma cell” to “Plasma_cells”; “Mast cell” to “Mast_cells”,

### Single cell or nuclei RNA-sequencing analysis

#### Normalization and highly variable gene detection

We normalized the total UMIs per nucleus to 10,000 and log-transformed these values using the LogNormalize() function in Seurat. The top 2000 genes with highly variable expression were identified using the “vst” method and subsequently, expression values were scaled across all samples and cells in a given dataset. Scaled *z* score residuals (‘relative expression’) were used for dimensionality reduction.

#### Dimensionality reduction and clustering

Principal component analysis (PCA) was conducted on the top 2000 highly variable genes as implemented in Seurat (RunPCA() function). Significant principal components were determined by the inflection point in a scree plot and used as input for non-linear dimensionality reduction techniques and batch-correction methods as described below. Uniform manifold approximation and projection (UMAP) was performed on significant PCs with 30 nearest neighbors for visualization in two dimensions. When required, we clustered datasets with the Seurat implementation of the leiden algorithm with a resolution of 1.5. This workflow corresponds to the “Uncorrected” analysis for each dataset.

#### Cell annotation

For the glioma snRNA-seq dataset (Richards-GBM-LGG), we annotated clusters with SingleR and a custom glioma scRNA-seq reference made up of 5 public datasets^10,15,36–38^. For all public datasets, we used published cell annotations provided by the authors. Some cell labels were adjusted for consistency across datasets as described above.

### Data integration

#### fastMNN

Fast matching mutual nearest neighbors for batch correction (fastMNN) algorithm identifies mutual nearest neighbours (MNNs) between datasets in the PCA reduced dimension space^21^. The resulting MNNs are used to align datasets into a shared space. fastMNN analysis was performed in the R programming environment (v.3.6.1). We used the Seurat (v.3.2.0) pre-processing workflow to normalize data, scale data, identify the top 2000 variable genes and run PCA as described above before integration. We used the SeuratWrappers (v.0.2.0) and batchelor (v.1.2.4) package to run fastMNN on variable genes. The top 20 resulting MNNs were then used for downstream UMAP reduction.

#### Conos

Clustering On Network of Samples^20^ (Conos) uses a joint graph to identify and co-localize subpopulations across different samples. First, we applied a standard Seurat (v.4.0.2) pre-processing workflow to normalize data, identify the top 2000 highly variable genes, scale data and run PCA on each batch (sample) individually in the R programming environment (v.4.0.0). We used the conos (v.1.4.1) and SeuratWrappers (v.0.3.0) R packages to perform pairwise comparisons between datasets to identify inter-sample mappings. Then, we used these inter-sample edges to construct a joint graph using the first 30 PCs, the top 2000 overdispersed genes, an inter-sample neighbour size (k) of 15, an within-sample neighbourhood size (k.self) of 5 and mutual nearest neighbours matching method. The joint graph was then embedded and visualized with LargeVis.

#### Harmony

Harmony is an unsupervised joint embedding method that employs an iterative clustering approach to align cells from different batches^23^. First the algorithm, embeds cells into reduced PCA space and soft assigns cells to clusters with favour towards mixed dataset representation. Then Harmony calculates both cluster and dataset centroids, and uses these centroids to calculate a correction factor for each dataset. The correction factor is then used to correct each cell with a cell-specific factor to move cells based on soft cluster membership and improve batch mixing. This process is repeated until convergence. In this study, we used the Seurat (v.3.2.0) pre-processing workflow to normalize data, scale data, identify the top 2000 variable genes and run PCA as described above before integration. We executed the Harmony algorithm through the harmony (v.1.0) and Seurat Wrappers (v.0.2.0) packages in the R programming environment (v.3.6.1). We ran Harmony using 50 PCs and a default theta value of 2 and lambda value of 1. The top 20 resulting harmony factors were then used for downstream UMAP reduction.

#### RPCA in Seurat

The Seurat implementation of reciprocal principal component analysis^26^ (RPCA) employs a similar methodology to canonical correlation analysis^19^ (CCA), where anchors are determined between datasets using RPCA and then each dataset is projected into the others PCA space with mutual neighbourhood constraints on the anchors to harmonize datasets. CCA identifies shared sources of variation and works well on very similar datasets with conserved anchors, but is not ideal for datasets of mixed composition such as malignant samples. For this reason, we chose to only benchmark RPCA which is a more conservative approach and less likely to overcorrect different biological cell states present between batches (samples) which are common between tumors. We used the Seurat (v.4.0.2) pre-processing workflow to normalize data and identify the top 200 highly variable genes individually within each batch (sample) in the R programming environment (v.4.0.0). We then selected the top 2000 genes that are repeatedly variable across batches as the integration features. Integration features were used for data scaling and PCA of each sample individually. We identified shared anchors across datasets using reciprocal PCA and a k.anchor value of 5. Finally, these anchors were used to integrate data using a k.weight of 100. If data integration was unsuccessful, we systematically reduced the k.weight by increments of 10. All datasets had a k.weight of 100, with the exception of Ma-LIHC (k.weight=50) and Yost-BCC (k.weight=80). We then ran data scaling, PCA and UMAP reduction using the top 20 PCs on the integrated dataset.

#### STACAS

Sub-Type Anchor Correction for Alignment in Seurat^24^ (STACAS) is an algorithm, similar to RPCA and CCA, that uses reciprocal PCA to identify anchors, project datasets into a shared reduced PCA space and then calculate mutation nearest neighbours. STACAS is designed for datasets that share only a subset of cells, as in tumor samples. Unlike RPCA and CCA, STACAS corrects batch effects while preserving biological heterogeneity by filtering aberrant integration anchors with a distance measurement and constructing an optimized sample-ordering tree for integration. Additionally, STACAS does not rescale gene expression to have a mean and variance of 0 before PCA, as this can mask biological signals between datasets. For each sample, we used Seurat (v.4.0.2) to normalize the data and identify the top 2000 highly variable genes, excluding mitochondrial and ribosomal genes. We then identified anchors between datasets using 2000 highly variable genes shared across batches, and filtered these anchors based on pairwise reciprocal PCA distance (dist.pct) of 0.8 using the STACAS (v.1.1.0) R package. Next, STACAS determined the optimal hierarchical tree for Seurat integration using filtered anchors and integrated the samples accordingly using the IntegrateData() function in Seurat with a k.weight of 100. If data integration was unsuccessful, we systematically tested k.weight and dist.pct combinations by reducing k.weight in increments of 10 and increasing dist.pct in increments of 0.05. Richards-GBM-LGG and Bi-RCC integrated with default parameters (k.weight=100, dist.pct=0.8), while Caron-ALL (k.weight=100, dist.pct=0.95) and Ma-LIHC (k.weight=20, dist.pct=0.8) required parameter optimization. Despite attempting >45 k.weight and dist.pct combinations, we were unable to generate a STACAS integration for Yost-BCC, perhaps because of the low cell count for some samples in the dataset. Integrated data was then scaled and reduced using PCA and UMAP on the top 20 PCs.

### Calculation of evaluation metrics

#### Local Inverse Simpson’s Index (LISI) metric

We used LISI scores to evaluate mixing between batches (samples) across entire datasets and within individual cell types. We calculated LISI scores using the compute_lisi() function in the immunogenomics R package (https://github.com/immunogenomics/LISI; v.1.0)^23^ with a perplexity of 30. If a cell type had less than 40 cells, we adjusted the perplexity to 10. We used UMAP (Uncorrected, RPCA, Harmony, STACAS, fastMNN) or LargeVis (Conos) cell embeddings as input for LISI scores. A high LISI score approaching the number of categorical variables present (i.e. batches or samples) indicates good mixing between groups in the local neighbourhood of a given cell. For example, the Bi-RCC dataset has 8 batches (samples), therefore an effective batch correction would have a LISI score around 8. To facilitate comparison of LISI scores across studies, we normalized the scores between [0,1] by dividing by the number of batches present within a given category. To accommodate the variable cellular composition between samples, we scaled by the number of batches with at least 1 cell present during cell-type specific LISI calculations. For visualizations, we calculated the median LISI score across cells for each categorical variable.

#### Runtime

We calculated the runtime of each method using the time function available in R (v.3.6.1 or v.4.0.0) environment. We did not time the pre-filtering steps (variable gene identification, PCA, etc.), and only measured runtime of the main integration function for each method.

### Statistics and reproducibility

No statistical method was used to predetermine sample size or cellular composition before data integration. Glioma nuclei with insufficient library complexity were excluded from the analyses as described in the methods. All plotting and statistical analysis was performed in the R statistical environment (v.3.6.1 and v.4.0.0), with the exception of doublet detection which was performed in a Python environment (v.3.6.1). We used the following plotting packages in R: ggplot2^39^ (v.3.3.3), ggpubr (v.0.4.0) (https://github.com/kassambara/ggpubr), ComplexHeatmap^40^ (v.2.5.4).

### Data sharing

To promote reproducibility and open-access, data generated and re-processed in this study has been shared through the interactive single-cell analysis portal CReSCENT^41^ (https://crescent.cloud/; Study IDs CRES-P24, CRES-P25, CRES-P26, CRES-P27, CRES-P28). For each study, we uploaded raw count and normalized expression matrices. For each integration method within a study, we uploaded UMAP coordinates **(Figures 1**, **S2**, **S3)** and cell-level metadata files detailing cell annotations, sample IDs, patient IDs and transcriptional clusters.

## DECLARATIONS

### Ethics approval and consent to participate

All samples were obtained following informed consent from patients.

### Consent for publication

Not applicable.

### Availability of data and material

The majority of data used during this study was obtained from publicly available sources. All datasets have been made available in their re-processed form through CReSCENT (https://crescent.cloud/; Study IDs CRES-P24, CRES-P25, CRES-P26, CRES-P27, CRES-P28). Code necessary to reproduce analyses and visualizations presented in this study are available without restrictions at https://github.com/pughlab/cancer-scrna-integration.

### Authors’ contributions

LMR performed all computational analysis, data curation, interpretation and visualization. MR generated single nuclei RNA-sequencing data for glioma samples. SM uploaded datasets to CReSCENT. SA, DCC and JJDM provided assistance with data interpretation. FJC and PBD provided tumor tissue for gliomas. LMR and TJP designed the study and wrote the manuscript. All authors read and approved the final manuscript.

## Acknowledgments

We thank the staff of the Princess Margaret Genomics Centre (www.pmgenomics.ca) for their expertise in generating the glioma single nuclei RNA-sequencing data used in this study.

## SUPPLEMENTARY FIGURE & TABLE LEGENDS

**Table S1. Clinical characteristics of glioma samples.** N.D. denotes no data available.

**Figure S1.**
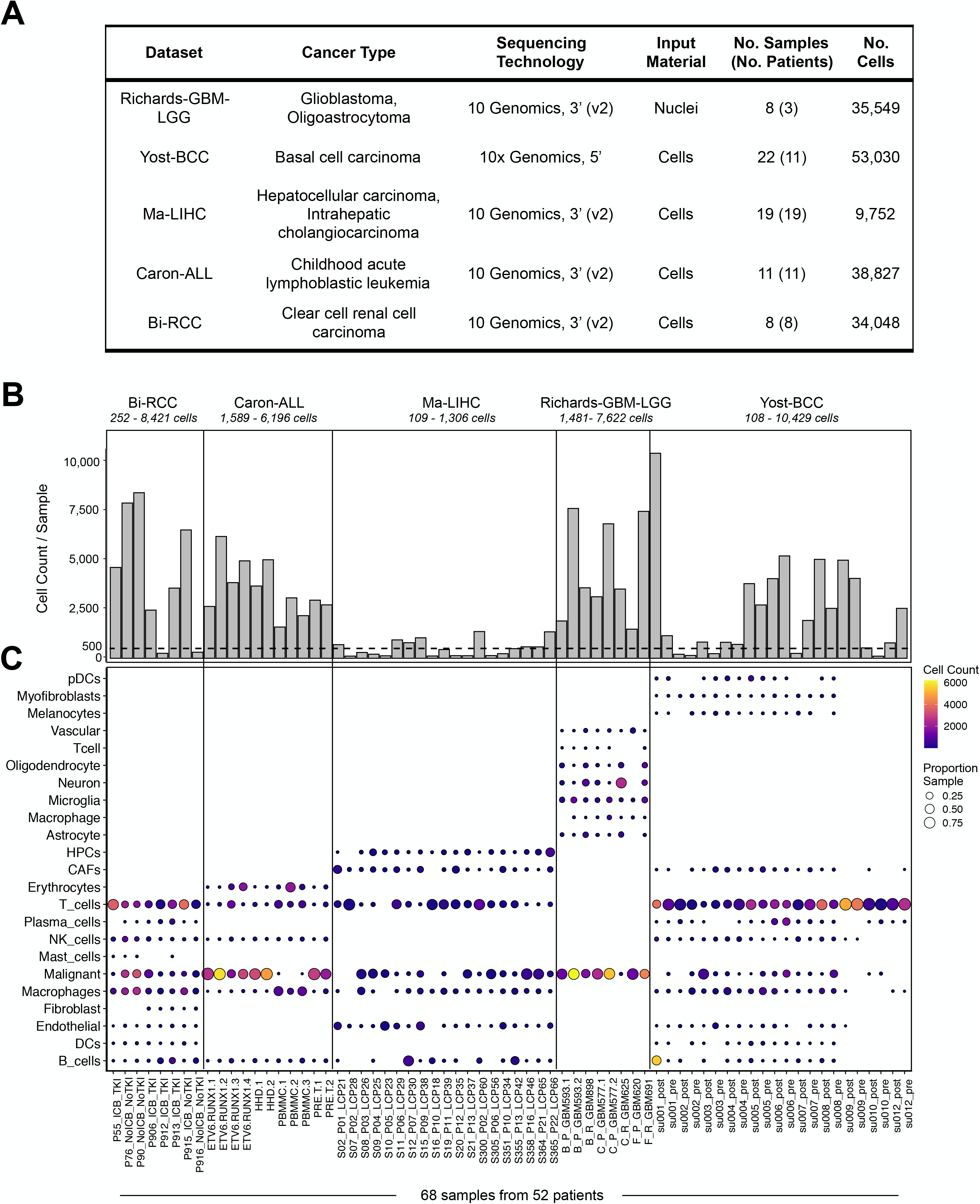
Cell type composition across datasets. **(A)** Table outlining cancer single-cell RNA sequencing datasets used in benchmark. **(B)** Histogram depicting the number of cells or nuclei profiled (y-axis) per sample (x-axis). Samples are ordered alphabetically within datasets, as shown in Panel C. Vertical solid lines separate samples from different datasets. Dashed horizontal line marks 500 cells. Values under study represent range across samples. **(C)** Dot plot representing the proportion and frequency of cell types (n=23; x-axis) across samples (n=68; y-axis). Dot size represents the portion of each sample corresponding to a given cell type. Dot color represents the discrete number of cells per sample for each cell type. Cell types with no dot are absent from a given sample. pDCs, plasmacytoid dendritic cells; DCs, dendritic cells; HPCs, hepatic precursor cells; CAFs, cancer associated fibroblasts.

**Figure S2.**
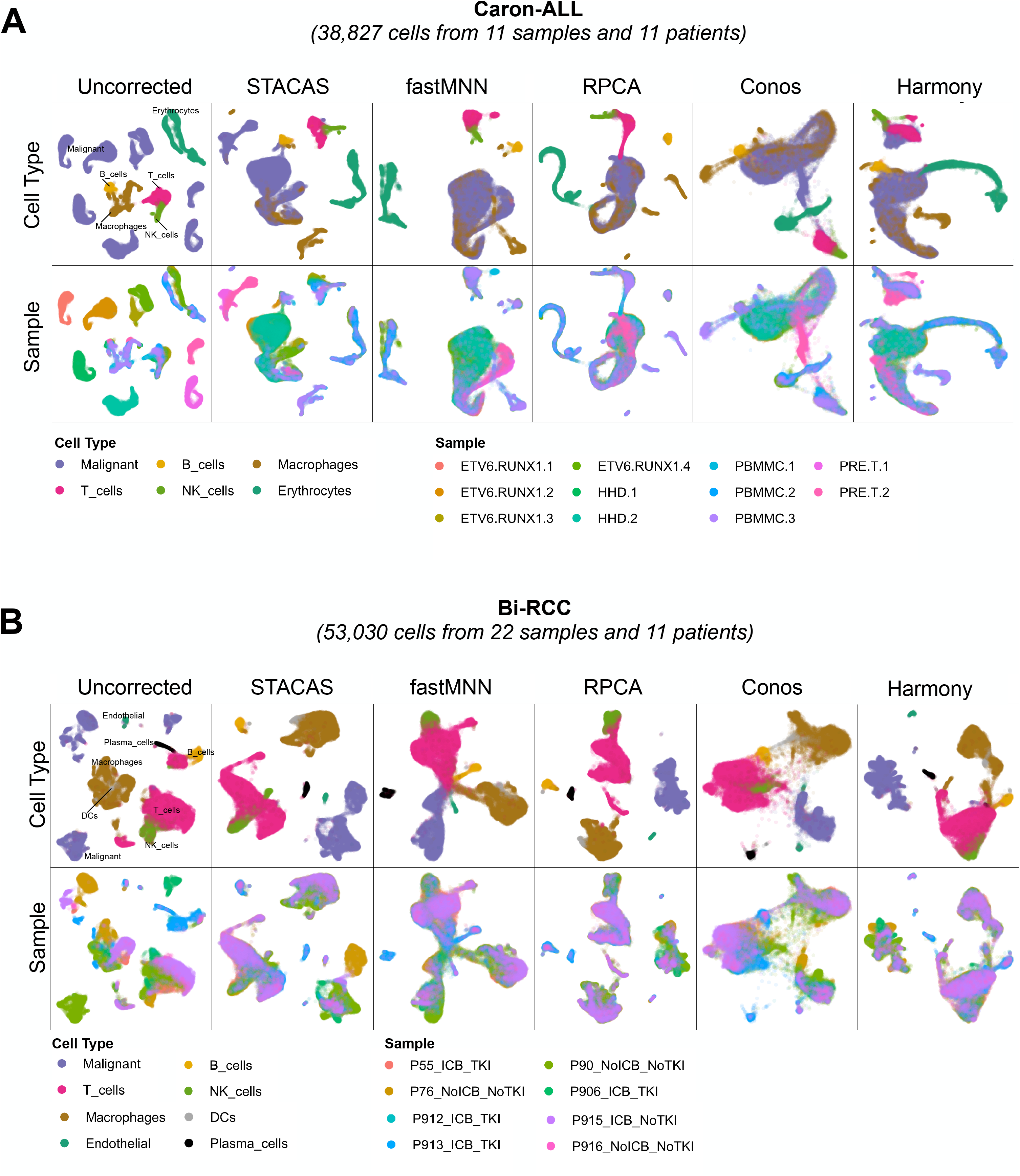
Comparison of data integration methods applied to childhood acute lymphoblastic leukemia and clear cell renal cell carcinoma datasets. UMAP or LargeVis (Conos) visualization of cells from **(A)** childhood acute lymphoblastic (Caron-ALL; n=38,827 cells) and **(B)** clear cell renal cell carcinoma (Bi-RCC; n=53,030 cells). Each dot represents a cell. Cells are colored by cell type (top row) or sample (bottom row). Columns represent either uncorrected (first column) or integrated data (columns 2 to 6). DCs, dendritic cells.

**Figure S3.**
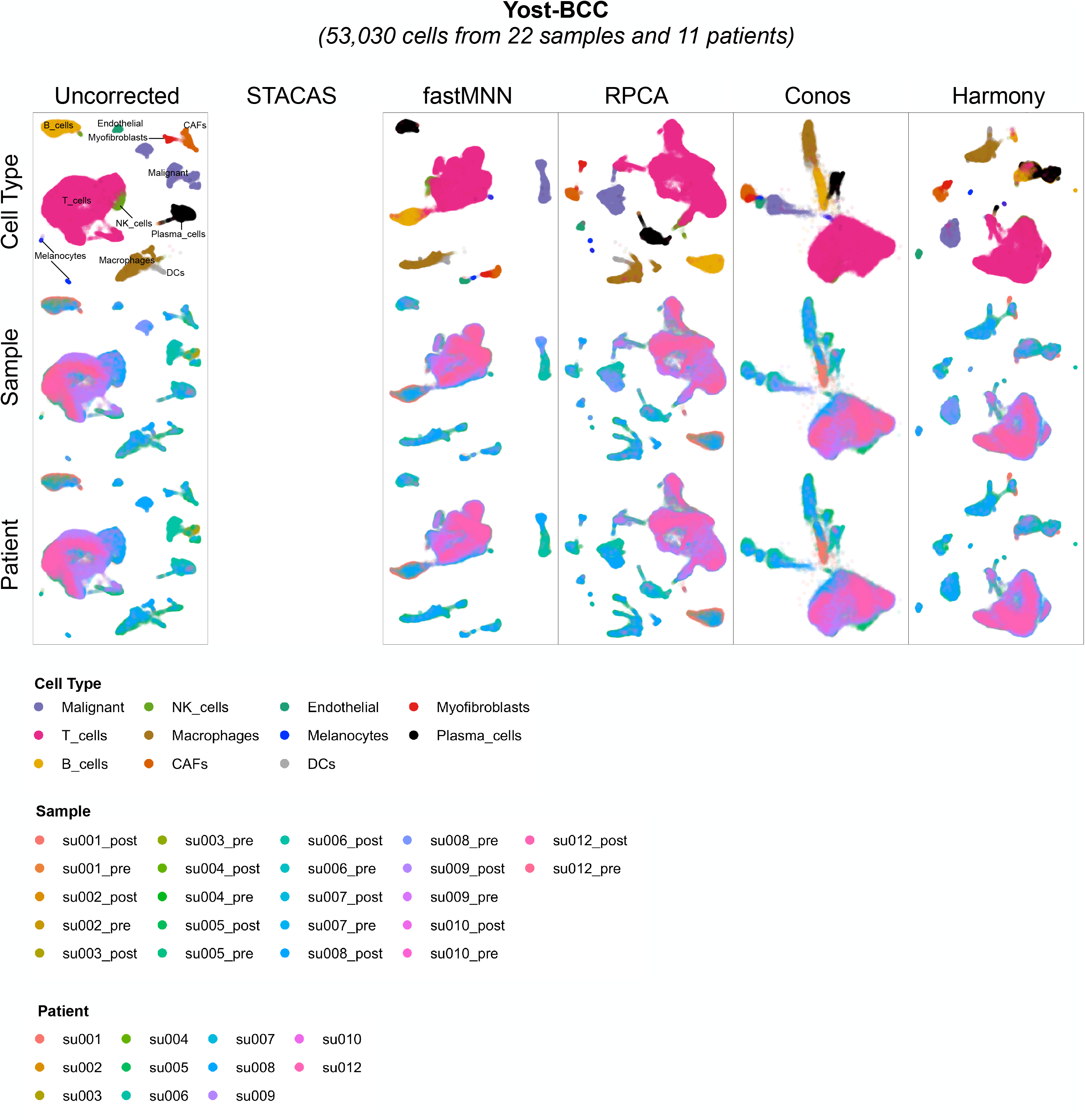
Comparison of data integration methods applied to basal cell carcinoma dataset. UMAP or LargeVis (Conos) visualization of cells from basal cell carcinoma (Yost-BCC; n=53,030 cells). Each dot represents a cell. Cells are colored by cell type (top row), sample (middle row) or patient (bottom row). Columns represent either uncorrected (first column) or integrated data (columns 2 to 6). DCs, dendritic cells; CAFs, cancer associated fibroblasts.

**Figure S4.**
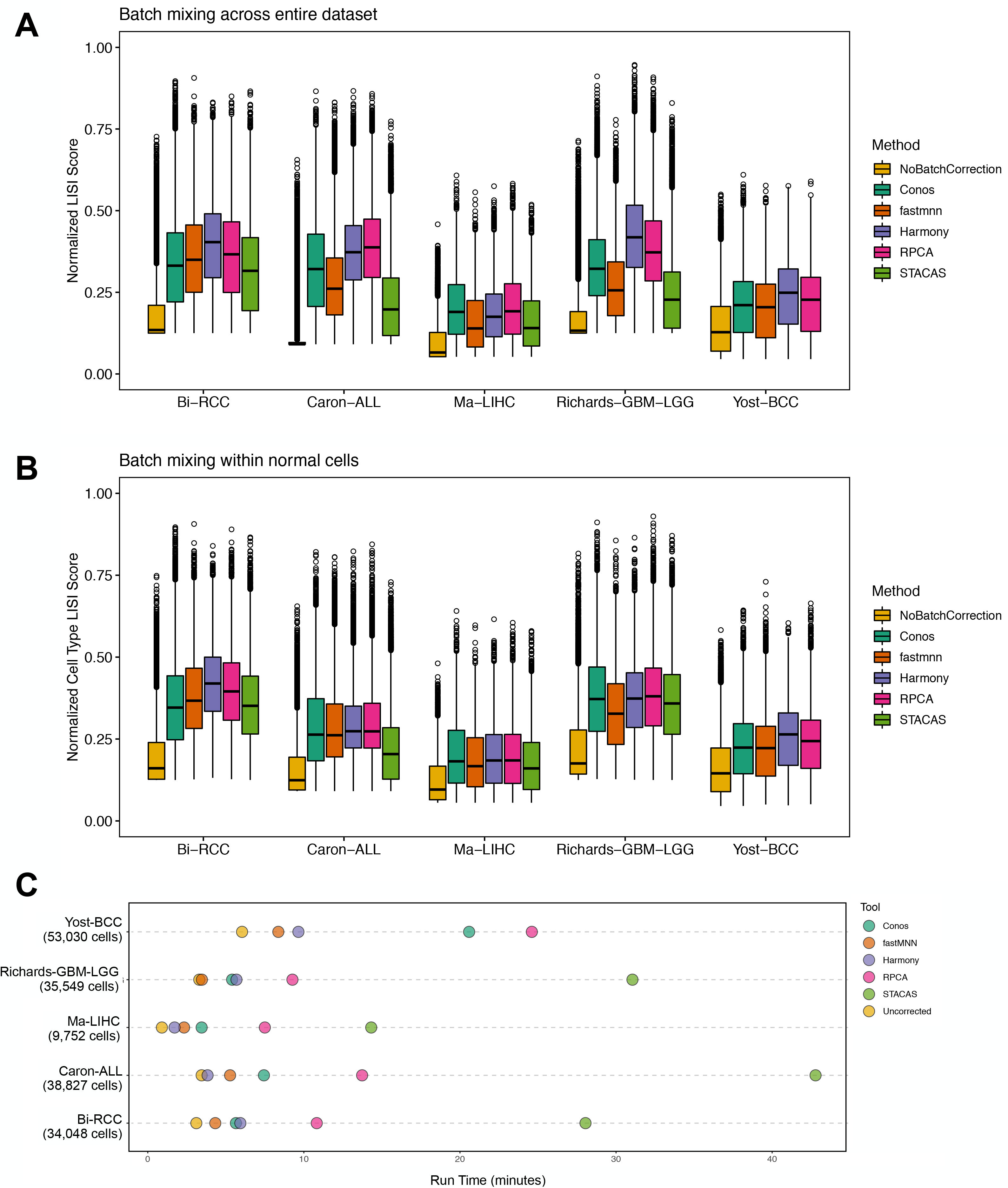
Evaluation of data integration performance and run time. Box plots grouped by dataset (x-axis) comparing normalized LISI score (y-axis) distributions between methods. Box plots are colored by data integration method. Box plots represent the median, first and third quartiles of the distribution and whiskers represent either 1.5-times interquartile range or most extreme value. Outliers are represented by open circles. **(A)** LISI scores represent sample mixing across entire datasets. **(B)** LISI scores represent sample mixing within normal cell types, excluding malignant cells. **(C)** Dot plot depicting run time in minutes (x-axis) for data integration method across datasets (y-axis). Dots are colored by data integration method.

## REFERENCES

1. Hicks, S. C., Townes, F. W., Teng, M. & Irizarry, R. A. Missing data and technical variability in single-cell RNA-sequencing experiments. Biostatistics 19, 562–578 (2018).

2. Slyper, M. et al. A single-cell and single-nucleus RNA-Seq toolbox for fresh and frozen human tumors. Nature Medicine 26, 792–802 (2020).

3. Haque, A., Engel, J., Teichmann, S. A. & Lönnberg, T. A practical guide to single-cell RNA-sequencing for biomedical research and clinical applications. Genome Medicine 9, 75 (2017).

4. Tran, H. T. N. et al. A benchmark of batch-effect correction methods for single-cell RNA sequencing data. Genome Biology 21, 12 (2020).

5. Luecken, M. D. et al. Benchmarking atlas-level data integration in single-cell genomics. bioRxiv 2020.05.22.111161 (2020) doi:10.1101/2020.05.22.111161.

6. Lütge, A. et al. CellMixS: quantifying and visualizing batch effects in single-cell RNA-seq data. Life Science Alliance 4, (2021).

7. Zanini, F. et al. Northstar enables automatic classification of known and novel cell types from tumor samples. Scientific Reports 10, 15251 (2020).

8. Tirosh, I. et al. Dissecting the multicellular ecosystem of metastatic melanoma by single-cell RNA-seq. Science 352, 189–196 (2016).

9. Puram, S. V. et al. Single-Cell Transcriptomic Analysis of Primary and Metastatic Tumor Ecosystems in Head and Neck Cancer. Cell 171, 1611–1624.e24 (2017).

10. Tirosh, I. et al. Single-cell RNA-seq supports a developmental hierarchy in human oligodendroglioma. Nature 539, 309–313 (2016).

11. Richards, L. M. et al. Gradient of Developmental and Injury Response transcriptional states defines functional vulnerabilities underpinning glioblastoma heterogeneity. Nature Cancer 2, 157–173 (2021).

12. Kinker, G. S. et al. Pan-cancer single-cell RNA-seq identifies recurring programs of cellular heterogeneity. Nature Genetics 52, 1208–1218 (2020).

13. Gojo, J. et al. Single-Cell RNA-Seq Reveals Cellular Hierarchies and Impaired Developmental Trajectories in Pediatric Ependymoma. Cancer Cell 38, 44–59.e9 (2020).

14. Bhaduri, A. et al. Outer Radial Glia-like Cancer Stem Cells Contribute to Heterogeneity of Glioblastoma. Cell Stem Cell 26, 48–63.e6 (2020).

15. Neftel, C. et al. An Integrative Model of Cellular States, Plasticity, and Genetics for Glioblastoma. Cell (2019) doi:10.1016/j.cell.2019.06.024.

16. Laughney, A. M. et al. Regenerative lineages and immune-mediated pruning in lung cancer metastasis. Nature Medicine 1–11 (2020) doi:10.1038/s41591-019-0750-6.

17. van Galen, P. et al. Single-Cell RNA-Seq Reveals AML Hierarchies Relevant to Disease Progression and Immunity. Cell 176, 1265–1281.e24 (2019).

18. Butler, A., Hoffman, P., Smibert, P., Papalexi, E. & Satija, R. Integrating single-cell transcriptomic data across different conditions, technologies, and species. Nature Biotechnology 36, 411–420 (2018).

19. Stuart, T. et al. Comprehensive Integration of Single-Cell Data. Cell 177, 1888–1902.e21 (2019).

20. Barkas, N. et al. Joint analysis of heterogeneous single-cell RNA-seq dataset collections. Nature Methods 16, 695–698 (2019).

21. Haghverdi, L., Lun, A. T. L., Morgan, M. D. & Marioni, J. C. Batch effects in single-cell RNA-sequencing data are corrected by matching mutual nearest neighbors. Nature Biotechnology 36, 421–427 (2018).

22. A description of the theory behind the fastMNN algorithm. https://marionilab.github.io/FurtherMNN2018/theory/description.html.

23. Korsunsky, I. et al. Fast, sensitive and accurate integration of single-cell data with Harmony. Nat Methods 16, 1289–1296 (2019).

24. Andreatta, M. & Carmona, S. J. STACAS: Sub-Type Anchor Correction for Alignment in Seurat to integrate single-cell RNA-seq data. Bioinformatics 37, 882–884 (2021).

25. Andreatta, M. et al. Interpretation of T cell states from single-cell transcriptomics data using reference atlases. Nat Commun 12, 2965 (2021).

26. Hao, Y. et al. Integrated analysis of multimodal single-cell data. Cell (2021) doi:10.1016/j.cell.2021.04.048.

27. Yost, K. E. et al. Clonal replacement of tumor-specific T cells following PD-1 blockade. Nature Medicine 25, 1251–1259 (2019).

28. Caron, M. et al. Single-cell analysis of childhood leukemia reveals a link between developmental states and ribosomal protein expression as a source of intra-individual heterogeneity. Sci Rep 10, 8079 (2020).

29. Ma, L. et al. Tumor Cell Biodiversity Drives Microenvironmental Reprogramming in Liver Cancer. Cancer Cell 0, (2019).

30. Bi, K. et al. Tumor and immune reprogramming during immunotherapy in advanced renal cell carcinoma. Cancer Cell 39, 649–661.e5 (2021).

31. Büttner, M., Miao, Z., Wolf, F. A., Teichmann, S. A. & Theis, F. J. A test metric for assessing single-cell RNA-seq batch correction. Nature Methods 16, 43 (2019).

32. Venteicher, A. S. et al. Decoupling genetics, lineages, and microenvironment in IDH-mutant gliomas by single-cell RNA-seq. Science 355, eaai8478 (2017).

33. Macosko, E. Z. et al. Highly Parallel Genome-wide Expression Profiling of Individual Cells Using Nanoliter Droplets. Cell 161, 1202–1214 (2015).

34. Alles, J. et al. Cell fixation and preservation for droplet-based single-cell transcriptomics. BMC Biology 15, 44 (2017).

35. Wolock, S. L., Lopez, R. & Klein, A. M. Scrublet: Computational Identification of Cell Doublets in Single-Cell Transcriptomic Data. Cell Systems 8, 281–291.e9 (2019).

36. Filbin, M. G. et al. Developmental and oncogenic programs in H3K27M gliomas dissected by single-cell RNA-seq. Science 360, 331–335 (2018).

37. Reitman, Z. J. et al. Mitogenic and progenitor gene programmes in single pilocytic astrocytoma cells. Nature Communications 10, 1–17 (2019).

38. Darmanis, S. et al. Single-Cell RNA-Seq Analysis of Infiltrating Neoplastic Cells at the Migrating Front of Human Glioblastoma. Cell Reports 21, 1399–1410 (2017).

39. Wickham, H. ggplot2: Elegant Graphics for Data Analysis. (Springer-Verlag, 2009). doi:10.1007/978-0-387-98141-3.

40. Gu, Z., Eils, R. & Schlesner, M. Complex heatmaps reveal patterns and correlations in multidimensional genomic data. Bioinformatics 32, 2847–2849 (2016).

41. Mohanraj, S. et al. CReSCENT: CanceR Single Cell ExpressioN Toolkit. Nucleic Acids Res 48, W372–W379 (2020).

